# Analysis of PPI networks of transcriptomic expression identifies hub genes associated with Newcastle disease virus persistent infection in bladder cancer

**DOI:** 10.1101/2020.12.18.423493

**Authors:** Umar Ahmad, De Ming Chau, Suet Lin Chia, Khatijah Yusoff, Syahril Abdullah, Soon Choy Chan, Abhi Veerakumarasivam

**Author notes:** Correspondence to A.V.

## Abstract

**Motivation:** Bladder cancer cells acquire persistent infection associated with oncolytic Newcastle disease virus (NDV) in which its molecular events are still unclear. This poses a potential problem for oncolytic virus application for cancer therapy. To unravel the molecular mechanism underlying the development of NDV persistent infection in bladder cancer, we used mRNA expression profile of the persistently infected bladder cancer cells to construct PPI network.

**Results:** Based on path and module exploring in the PPI network, the bridges were found mainly from pathways of p53 signalling, ECM-receptor interaction, and TGF-beta signalling by the upregulated mRNAs, to the antigen processing and presentation, protein processing in endoplasmic reticulum, completement and coagulation cascades by the downregulated mRNAs in NDV persistent TCCSUPPi cells. In persistent EJ28Pi cells comparatively, connections were identified mainly from pathways of renal carcinoma, viral carcinogenesis, Ras signalling and cell cycle by the upregulated mRNAs, to the Wnt signalling, HTLV-I infection and pathways in cancer by the downregulated mRNAs. This connection was mainly dependent on of *RPL8- HSPA1A/HSPA4* in TCCSUPPi cells and *EP300, PTPN11, RAC1* - *TP53, SP1, CCND1* and *XPO1* in EJ28Pi cells. Oncomine validation showed that the top hub genes identified in the network that includes *RPL8, THBS1, F2* from TCCSUPPi and *TP53* and *RAC1* from EJ28Pi are involved in the development and progression of bladder cancer. Protein-drug interaction network, have identified several drugs targets that could be used to disconnect the linkages between modules and prevent bladder cancer cells from acquiring NDV persistent infection. This is the first time reporting the PPI network analysis of differentially expressed mRNAs of the NDV persistently infected bladder cancer cell lines which provide an insight into screening drugs that could be used together with NDV to manage bladder cancer resistance to therapy and progression.

## Introduction

Bladder cancer (BC) is a growth of abnormal tissue term a tumour that develops in the bladder epithelial lining and spread into the muscle layer in some cases[1]. It is also the tenth most common cancer in the world that account for 4.7% of all the new cancer cases[2]. BC poor prognosis is due to the limited treatment options for advance disease and resistance to conventional therapies. Newcastle disease virus (NDV) is one of the promising novel classes of cancer specific agents that kills human tumour cells while sparing normal cells unharmed[3–5]. This selective killing of cancer cells by NDV is due to the defects in antiviral responses such as the production of interferon which favours the viral replication[6]. The mechanism by which NDV kill cancer cells is via the activation of both intrinsic and extrinsic apoptosis pathways [7] and direct virus-mediated oncolysis[8]. Moreover, NDV triggers long-term adaptive immune-response against infected cancer cells [9], making it a strong agent for inducing cell death in cancer. However, NDV has been found to persistently infect subpopulation of cancer cells that resist NDV-mediated oncolysis [10]. Similar observations have been reported in other oncolytic viruses such as Reovirus [11] and Measles virus [12]. Persistent infection indicates the ability of a subpopulation of cancer cells to resist NDV-mediated oncolysis. Thus, persistent infection in cancer cells poses a potential problem for maximising the efficacy potential of oncolytic viruses in cancer therapy as tumour contains heterogenous subset of cells that harbour a spectrum of genetic aberrations. Although recombinants NDV could have enhanced oncolytic capability, the risk of persistent infection will still remain. The exact mechanism by which cancer cells acquire NDV persistent infection and the molecular basis underlying the development of persistent infection in BC has not been completely elucidated.

Thus, to unravel the regulatory mechanism of oncolytic NDV persistent infection in bladder cancer at molecular level, we previously compared the mRNA expression differences between normal and cancer of the human bladder by transcriptomics [13]. The data indicated that a total of 63 and 134 mRNAs were obtained from TCCSUPPi and EJ28Pi relative to their control respectively, 25 mRNAs were upregulated (log2 fold-change ≥ 0) and 38 mRNAs were downregulated (log2 fold-change ≤ 0) in TCCSUPPi cells. Whereas, 55 mRNAs were upregulated (log_2_ fold-change ≥ 0) and 79 mRNAs were downregulated (log_2_ fold-change ≤ 0) in EJ28Pi cells [13]. These differentially expressed genes (DEGs) were significantly enriched in some important upregulated pathways such as TGF-beta signaling, KRAS signaling up and interferon gamma response. Moreover, the study suggested that evasion of apoptosis is crucial in development of persistent infection in bladder cancer cells. The data further revealed that some essential molecular functions such as calcium binding (GO:0005509) and DNA-binding transcription repressor activity, RNA polymerase II-specific (GO:0001227) in TCCSUPPi and protein domain specific binding (GO:0019904) and RNA polymerase II regulatory region sequence-specific DNA binding (GO:0000977) in EJ28Pi were significantly enriched. These may cooperatively contribute to the development of NDV persistent infection in bladder cancer.

Proteins function collectively via protein-protein interactions (PPI) within the cell. This interaction is essential for most of the biochemical activities to achieve specific function in a living cell [14, 15] and also provide single protein with multiple functions[16, 17]. Thus, employing PPI methodologies to unravel the molecular mechanisms of biological processes draw an increasing attention in recent time[15, 18, 19]. To deeply understand the regulatory mechanism mechanisms underlying the state of many diseases, PPI networks are generally performed by analysing the DEGs obtained from those diseases [17, 20, 21]. However, there is currently no PPI network analysis for DEGs obtained from an established persistently infected bladder cancer cell. In this study, all the total DEGs obtained in TCCSUPPi (63) and EJ28Pi (134) were analysed separately to construct a PPI network to better understand and unravel the molecular mechanism underlying the development of NDV persistent infection in bladder cancer.

## Results

### PPI network of DEGs from TCCSUPPi

To deeply understand the regulatory mechanisms employed by bladder cancer cell lines in developing NDV persistent infection, differentially expressed genes (DEGs) obtained from both persistent TCCSUPPi and EJ28Pi cell lines were used to construct protein-protein interactions (PPI) network through STRING Interactome database[22]. All the total DEGs obtained in TCCSUPPi (63) and EJ28Pi (134) were analyzed separately for network interactions. As a result, 6 subnetworks that included a continent (subnetwork 1) and 5 islands (subnetwork 2-6) were generated in TCCSUPPi cells. Two subnetworks with highest scores were selected for further analysis. The subnetwork 1 contained 291 nodes, 309 edges and 12 seeds (Figure 1A) and subnetwork 2 had 34 nodes, 36 edges and 2 seeds (Figure 2B). Expressions and the degrees of connection between nodes were represented by their colours and areas respectively. Number of hub nodes in the entire network were further analyzed and the top 14 hub nodes were selected and graphically presented (Figure 3C). Twelve (12) hub nodes out of 14 were mostly from subnetwork 1 and two superfamily member of cadherin *(CDH2* and *CDH5)* were clustered in subnetwork 2(Figure 2B) with *CDH2* upregulated and *CDH5* downregulated.

**Figure 1:**
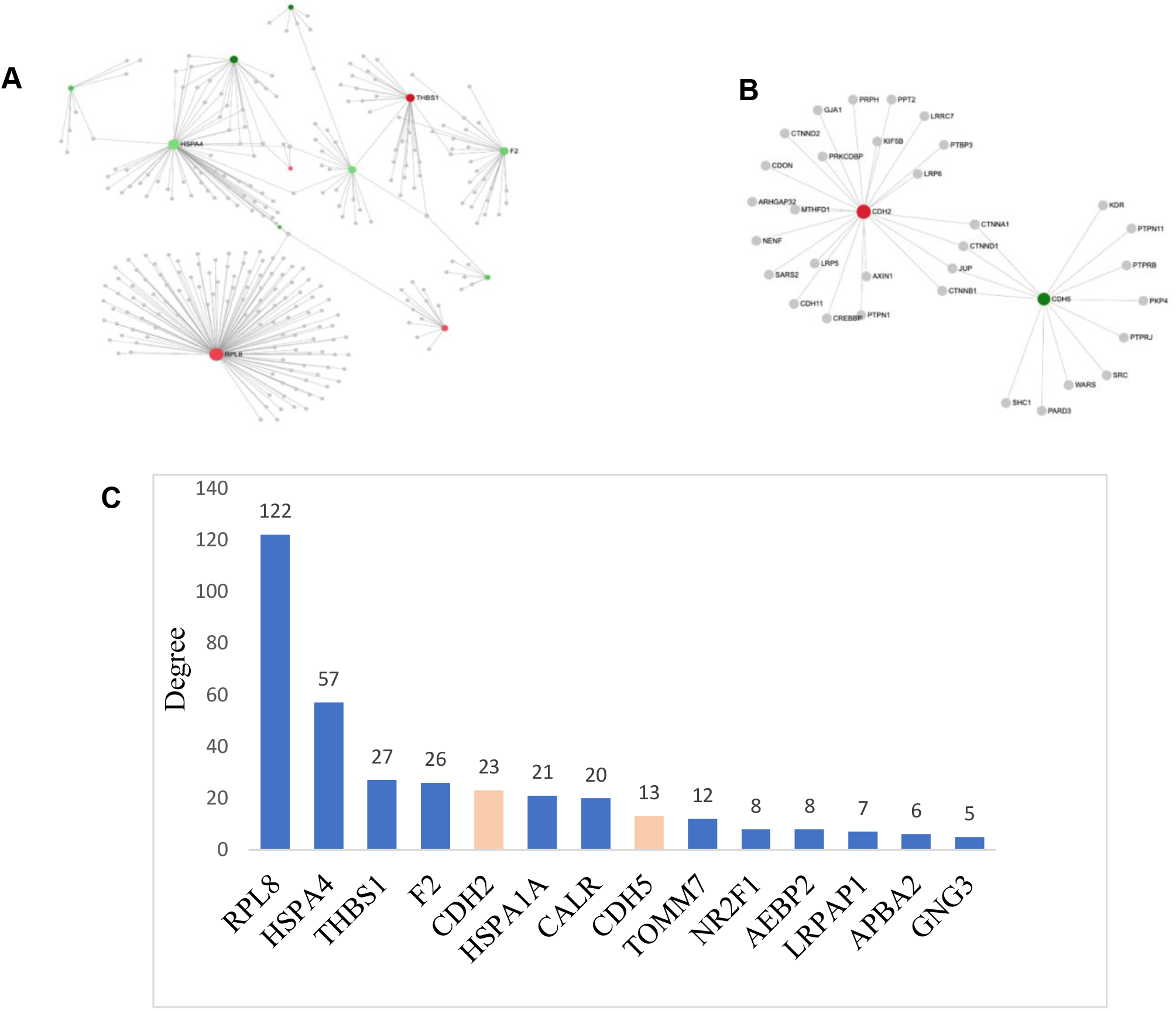
First and second identified networks in TCCSUPPi cells. A) First identified network in TCCSUPPi cells. Red and green colours represent the nodes expression that are up- and down-regulated, respectively. The expression level is represented by the shades of colour and the area of the nodes indicate the degrees that the nodes connect to others. Nodes with gene names are the top 4 nodes in the PPI network. B) Second identified network in TCCSUPPi cells. Subnetwork 2 contains both up- and downregulated nodes that are affected in the pathways. Nodes in red and green colours are upregulated and downregulated in TCCSUPPi cells, respectively. C) Hub nodes in the PPI network in TCCSUPPi cells. Top 14 hub nodes with their degree level are shown. Genes in blue colour are from subnetwork 1 and red accent are subnetwork 2.

### Functional connections in the network

Connections between functions in the identified network were further explored and related nodes were re-constructed (Figure 2). As illustrated, pathways of bladder cancer, malaria, mitophagy, p53 signaling, ECM-receptor interaction, TGF-beta signaling, phagosome, ribosome, focal adhesion and proteoglycans in cancer were significantly enriched (*p*<0.05) by the upregulated DEGs in the nodes connecting the PPI network (subnetwork 1) (Supplementary Table 1). Antigen processing and presentation, protein processing in endoplasmic reticulum, prion diseases, legionellosis, longevity regulating, complement and coagulation cascades, platelet activation and spliceosome pathways were significantly enriched (*p*<0.05) by the downregulated DEGs with connected nodes in the PPI network (subnetwork 1) (Supplementary Table 2).

To identify the nodes that are implicated in the aforementioned pathways, medullary analysis was carried out. A total of 9 functional clustered modules and related hub genes were discovered, however, only modules with majority of the nodes were used for redesigning of the modular network. Modules 0 and 1 were observed to contain significant number of the nodes (*p* < 0.05) that contributed to the activation of pathways mentioned above. The top two significant modules were presented in different colours (Figure 2). The results demonstrated that module 0 (coloured blue) and module 1 (coloured red) are key players in in the PPI network of the TCCSUPPi cells, which means that those clustered of genes in module 0 and module 1 act together to promote the development of NDV persistent infection in TCCSUP bladder cancer cell line. The results further illustrated how the upregulated *RPL8* group and downregulated *HSPA1A/HSPA4* are functionally connected (Figure 2).

**Figure 2:**
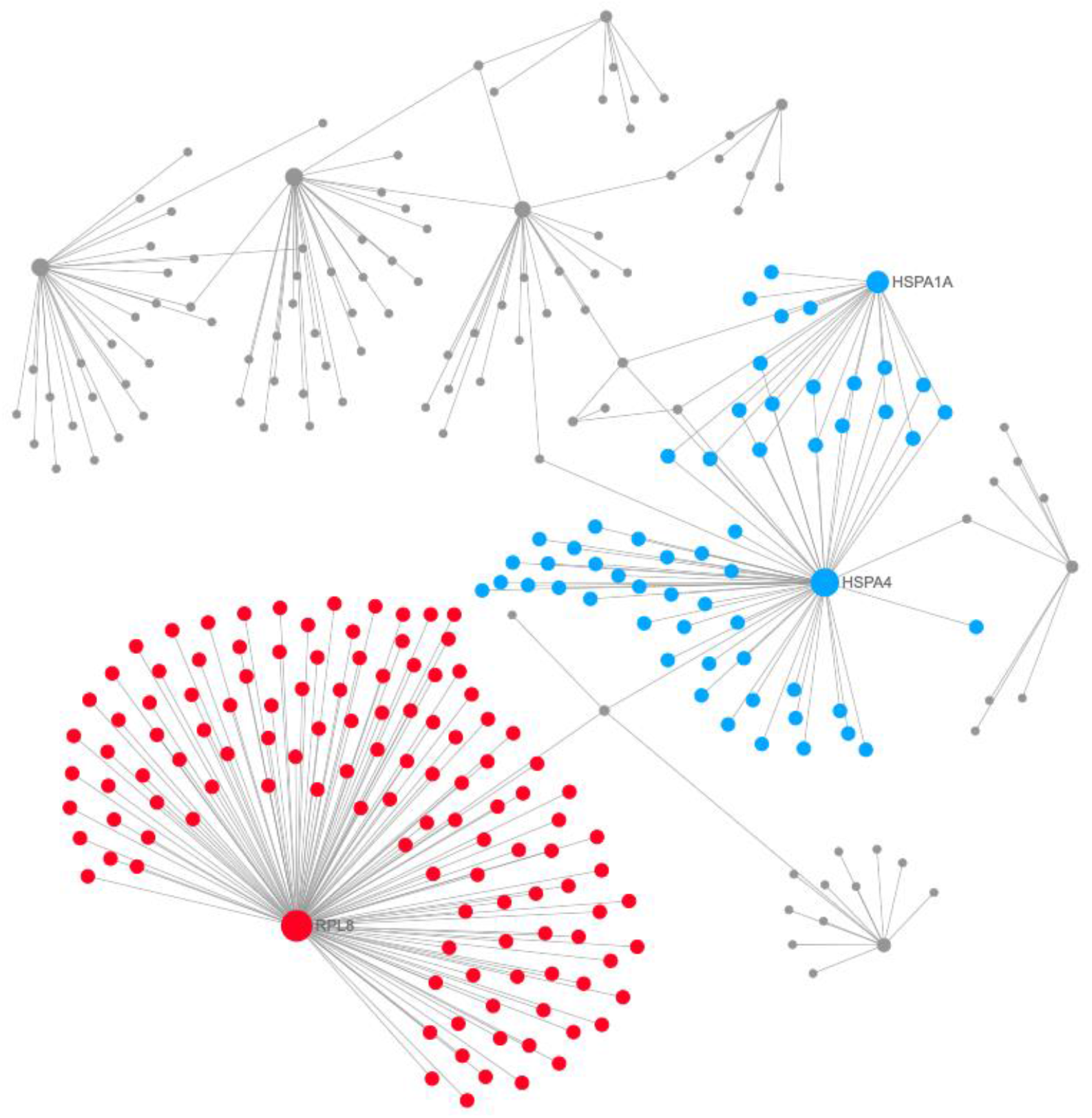
Modules 0 and 1 in the TCCSUPPi PPI network. The modules in blue and red are module 0 and module 1 respectively. The degrees of the nodes that connect to others in the network are represented by areas of the nodes.

### Protein drug interactions in TCCSUPPi

Moreover, to identify drug interactions between these connected nodes, we carried out protein-drug interaction analysis using the upregulated nodes that included *RPL8* and *THBS1* and downregulated nodes that included *F2* and *HSPA4.* Two subnetworks were identified. Subnetwork 1 comprises of 104 nodes, 103 edges and 1 seed while subnetwork 2 has 4 nodes, 3 edges and 1seed. Based on the analysis in subnetwork 1, several drugs were identified to be connected to the coagulation factor II, thrombin (*F2*) node. The top major drugs that are linked to *F2* are lepirudin, bivalirudin, drotrecogin alfa, coagulation factor IX (recombinant), menadione, argatroban, and proflavine (Figure 3A). The remaining list of drugs can be found in supplementary Table 3. In contrast to subnetwork 2, ribosomal protein L8 *(RPL8)* that was upregulated is linked to alpha-hydroxy-beta-phenyl-propionic acid, anisomycin and puromycin drugs (Figure 3B). These results demonstrate that the identified drugs above can be used to suppress both the upregulated and downregulated nodes, making it possible for the TCCSUP bladder cancer cells to prevent NDV persistent infection.

**Figure 3:**
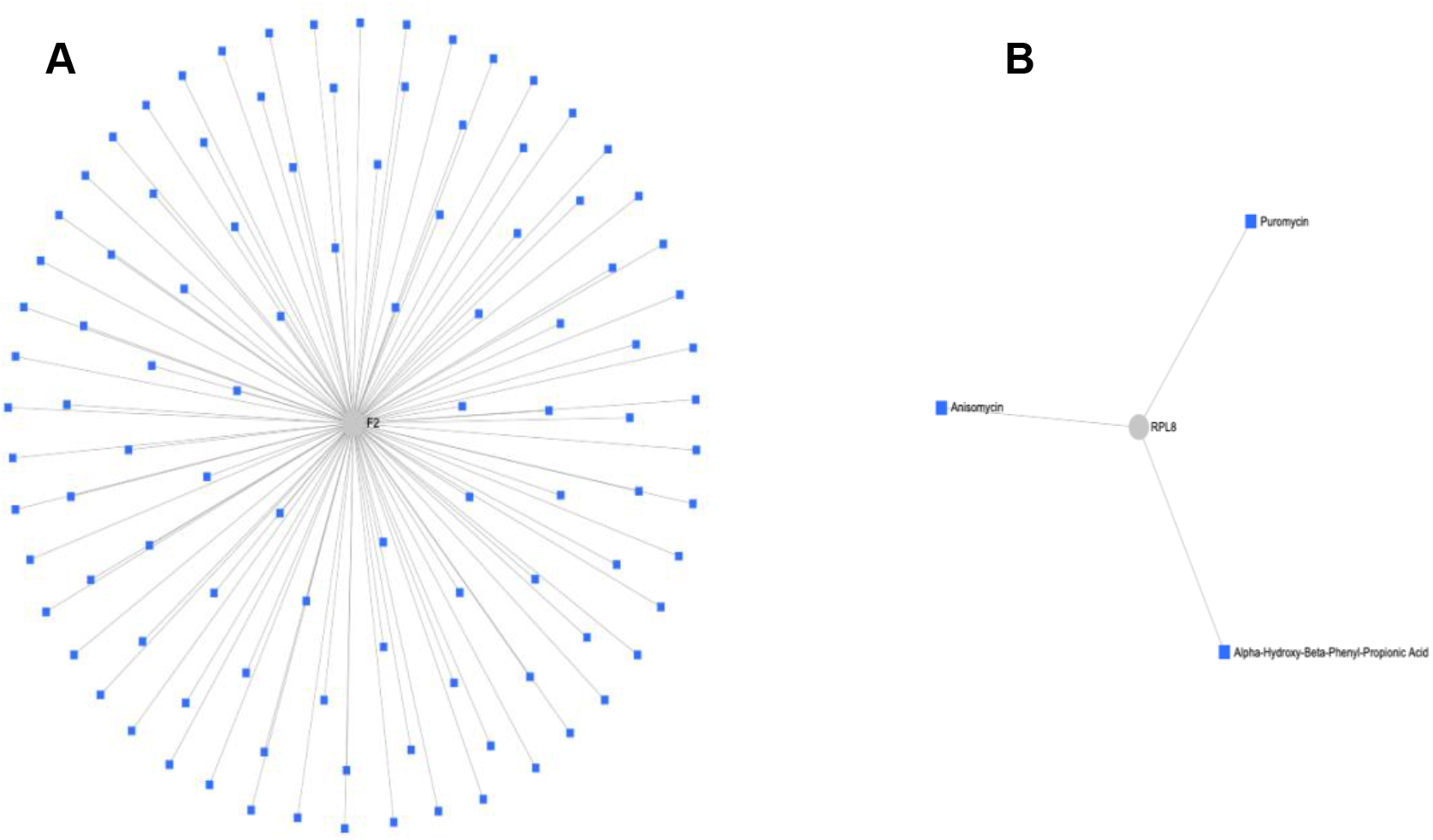
Protein-drug interaction network. A) The figure illustrates interactions between the downregulated node (*F2*) and several multiple drugs. B) The figure shows interactions between the upregulated node (*RPL8*) and three drugs.

### PPI network of DEGs from EJ28Pi

Subsequently, the 134 DEGs from EJ28Pi cells were analysed for protein-protein network interactions. The results display a network comprising of 14 subnetworks including one continent (subnetwork 1) and 13 islands (subnetwork 2 - 14). Once again, the network with the highest scores were selected and analysed in order to provide an insight into the mechanism employed by EJ28 cells in developing NDV persistent infection. Subnetwork 1 had 1161 nodes, 1662 edges and 57 seeds (Figure 4A) while subnetwork 2 identified 21 nodes, 20 edges and 1 seed (Figure 4B). Expression of each node is illustrated by their colours while degrees of connection between nodes were represented by areas. The top 16 hub nodes from the entire network analysis were assessed for their distribution and 15 hub nodes out of this total were from subnetwork 1 while only NADH ubiquinone oxidoreductase core subunit S2 *(NDUFS2)* was observed from subnetwork 2 (Figure 4C).

**Figure 4:**
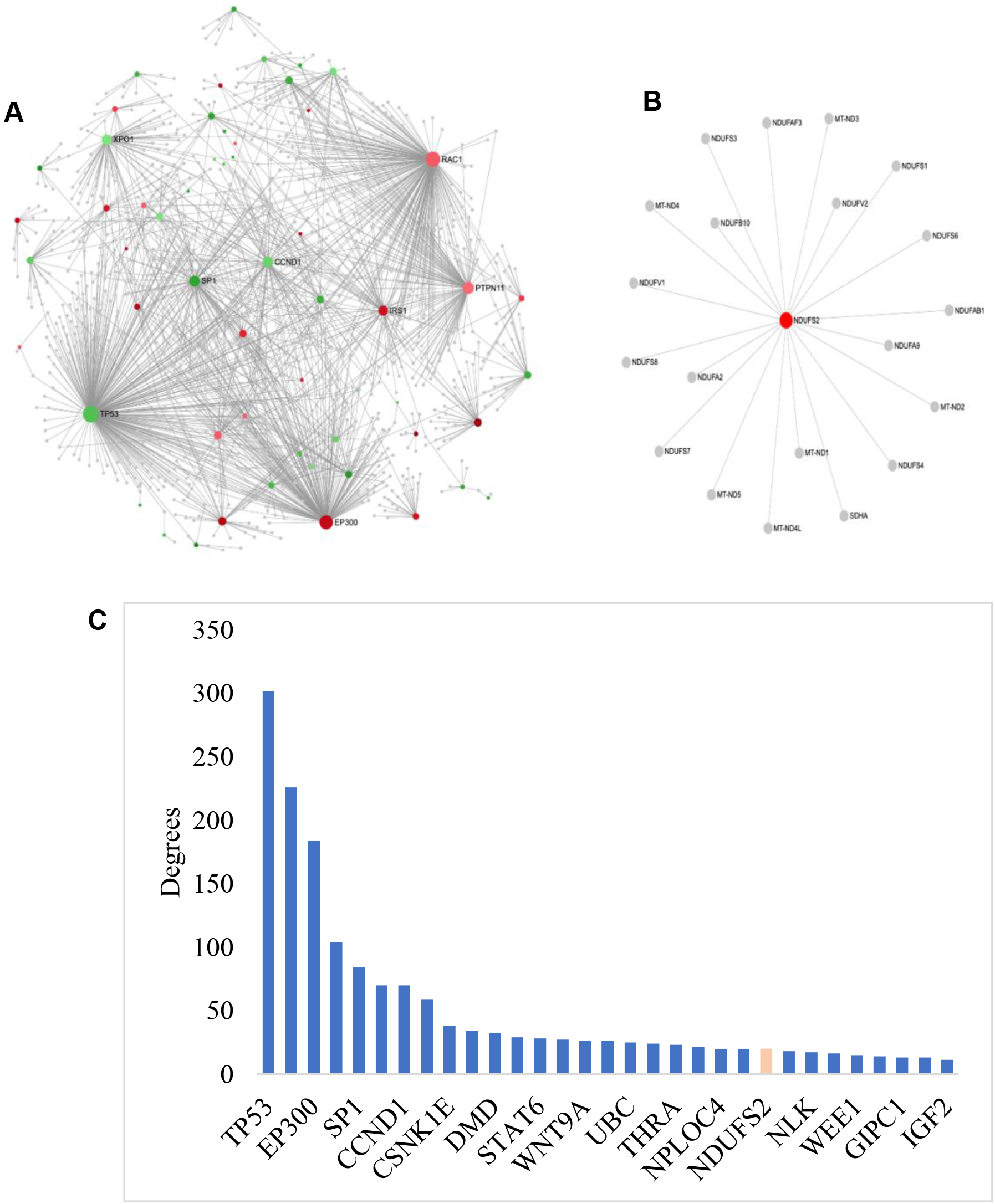
First and second identified networks in EJ28Pi cells. A) First identified network in EJ28Pi cells. Red and green colours represent the nodes expression that are up- and down-regulated. The expression level is represented by the colour grades and the area of the nodes indicate the degrees that the nodes connect to others. Nodes with gene names are the top 8 nodes in the PPI network. B) Second identified network in EJ28Pi cells. Subnetwork 2 showing only the upregulated node that is affected in the pathway. The red colour node is an indication that it is upregulated in EJ28Pi cells. C) Hub nodes in the PPI network in EJ28Pi cells. Top 16 hub nodes with their degree level are shown. Genes in blue colour are from subnetwork 1 and red accent are subnetwork 2.

Next, functional connections within the constructed network were studied and related nodes were re-designed (Figure 5). As shown, pathways of renal cell carcinoma, viral carcinogenesis, proteoglycans in cancer, prostate cancer, insulin resistance, Ras signalling, circadian rhythm, neurotrophin signalling, cell cycle, aldosterone-regulated sodium reabsorption, FoxO signalling, microRNAs in cancer, Wnt signalling, influenza A, tight junction, viral myocarditis, Kaposi’s sarcoma-associated herpesvirus infection, PI3K-Akt signalling, epithelial cell signaling in Helicobacter pylori infection, adipocytokine signalling, epstein-Barr virus infection, adherens junction, bacterial invasion of epithelial cells, cAMP signalling, HTLV-I infection, Longevity regulating pathway, glucagon signaling pathway, leukocyte transendothelial migration, AMPK signaling pathway, pathways in cancer, natural killer cell mediated cytotoxicity and measles were significantly enriched *(p* <0.05) by the nodes containing upregulated DEGs connecting the PPI network (subnetwork 1). The complete list of these pathways with their false discovery rate (FDR) are listed in Supplementary Table 4.

**Figure 5:**
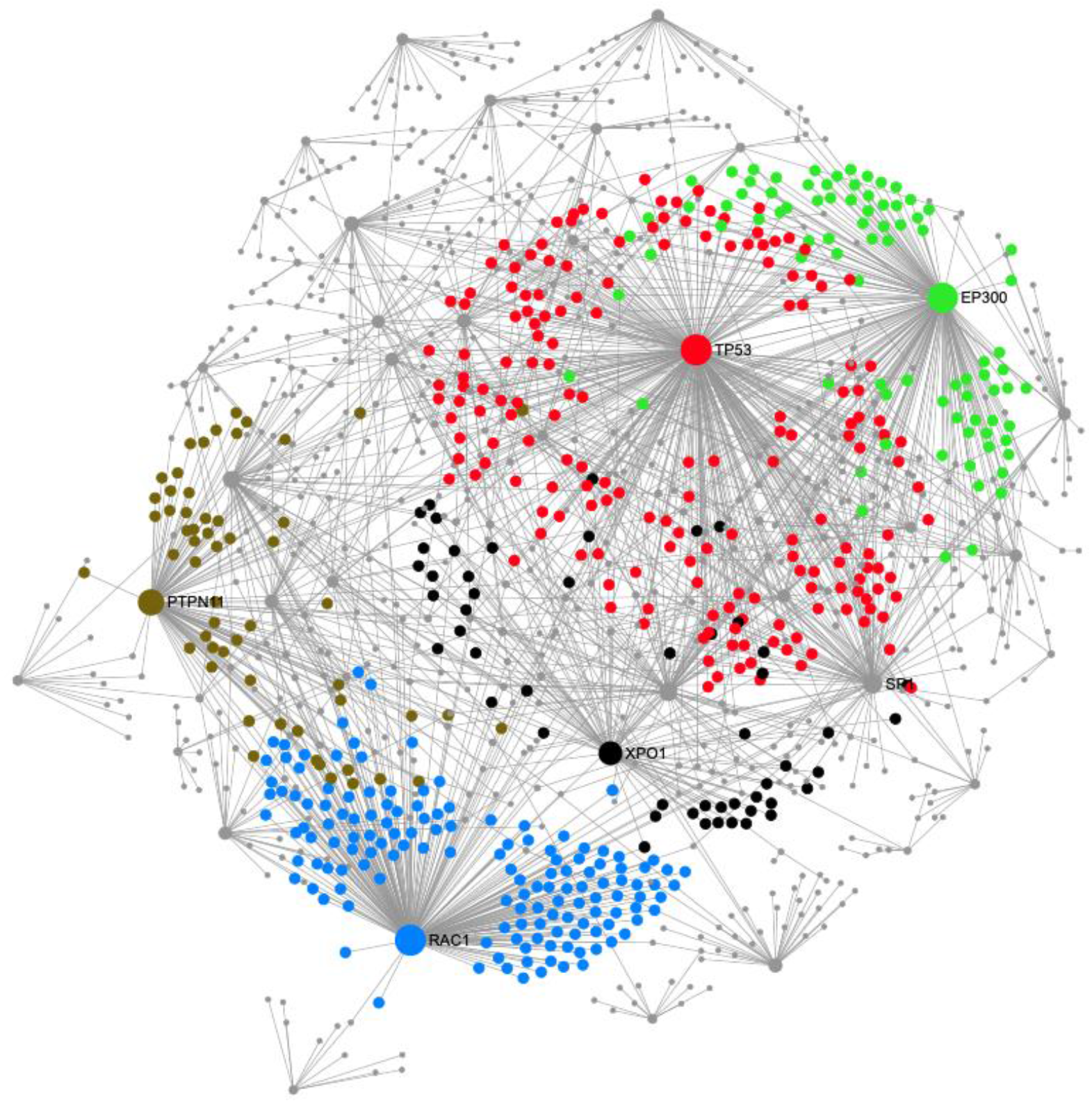
Modules 4, 5, 6, 7 and 8 in the EJ28Pi PPI network. The modules in red, blue, green, black and brown are for modules 4, 5, 6, 7, and 8 respectively. The degrees of the nodes that connect to others in the network is represented by areas of the nodes.

On the other hand, pathways of Wnt signaling, proteoglycans in cancer, HTLV-I infection, cancer, breast cancer, cellular senescence, melanoma, glioma, MAPK signaling, transcriptional misregulation in cancer, focal adhesion, prostate cancer, endocrine resistance, Th17 cell differentiation, thyroid hormone signaling, thyroid cancer, FoxO signaling, bladder cancer, measles, oxytocin signaling, hippo signaling, amyotrophic lateral sclerosis (ALS), hepatitis C, Jak-STAT signaling, hepatitis B, endometrial cancer, basal cell carcinoma, axon guidance, mitophagy - animal, central carbon metabolism in cancer, Inflammatory bowel disease (IBD), non-small cell lung cancer, Kaposi’s sarcoma-associated herpesvirus infection, long-term potentiation, amphetamine addiction, PI3K-Akt signaling, p53 signaling, platinum, drug resistance, pancreatic cancer, chronic myeloid leukemia, regulation of actin cytoskeleton, colorectal cancer, Th1 and Th2 cell differentiation, small cell lung cancer, choline metabolism in cancer, neurotrophin signaling pathway, cell cycle, platelet activation and oocyte meiosis were significantly enriched by the downregulated DEGs that have the connected nodes in the PPI network (subnetwork 1). The identified pathways with their false discovery rate (FDR) can be found in Supplementary Table 5.

As for the modules in the network, the PPI network consisted of 41 modules but only modules with majority of the nodes were selected for redesigning of the modular network. Modules 4, 5, 6, 7, and 8 had the most significant number of nodes (*p*<0.05) that contributed to the activation and enrichment of the above pathways. The modules are coloured red (module 4), blue (module 5), green (module 6), black (module 7) and brown (module 8) respectively (Figure 5). These five modules have significantly *(p* <0.05) act together to contribute to the development of NDV persistent infection in EJ28 bladder cancer cell line being that they functionally connected via upregulated *EP300, IRS1, PTPN11,* and *RAC1* groups as well as through downregulated *TP53, SP1, CCND1* and *XPO1* respectively.

To disconnect the linkage between these major modules in the PPI network (subnetwork 1), protein-drug interaction network analysis was performed. The upregulated nodes that included *EP300*, *IRS1*, *PTPN11*, and *RAC1,* as well as the downregulated nodes comprising of *TP53*, *SP1*, *CCND1* and *XPO1* were mapped to DrugBank database for matching nodes to obtain their drug interaction information. Two subnetworks were discovered with subnetwork 1 containing 4 nodes, 3 edges and 1 seed and subnetwork 2 containing 3 nodes, 2 edges and 1seed. Dextromethorphan and guanosine-5’-diphosphate drugs were identified to effectively interact with upregulated rac family small GTPase 1 (*RAC1*) node (Figure 6A). Contrary to subnetwork 2, tumour protein P53 *(TP53)* node that was downregulated is connected to acetylsalicylic acid, AZD, and 1-(9-ethyl-9H-carbazol-3-yl)-N-methylmethanamine drugs (Figure 6B), which basically means that these drugs can be used together with NDV to enhance the oncolytic activity of NDV against EJ28 bladder cancer cell lines and prevent the cells from acquiring persistent infection.

**Figure 6:**
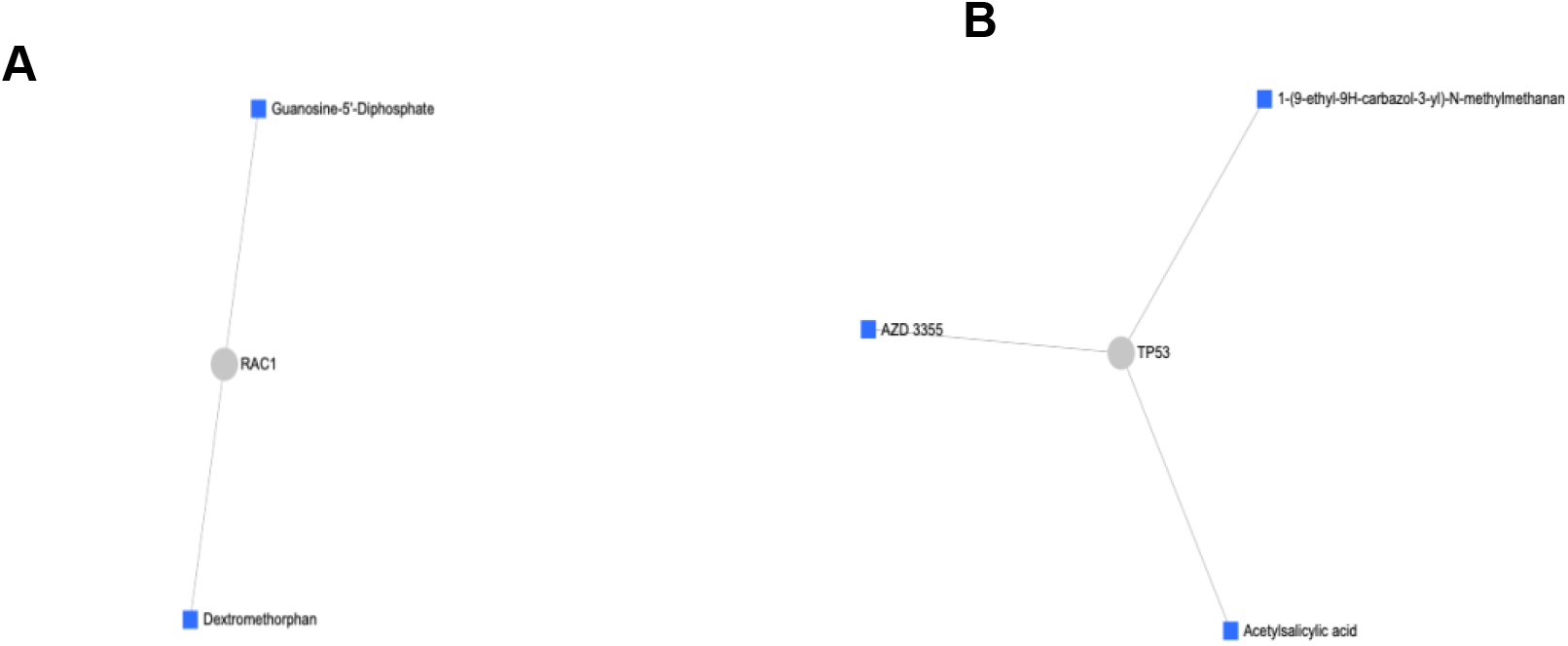
Protein-drug interaction network. A) The figure illustrates the interactions between upregulated node (*RAC1*) and two drugs. B) The figure shows interaction between upregulated node (*TP53*) and three drugs.

### Validation of hub genes in Oncomine

To validate the expression profiles of the top hub genes that were identified in the proteinprotein interaction network, mRNA expression mining of these hub genes in publicly available Oncomine database (www.oncomine.org) [23] that include the upregulated nodes *RPL8* and *THBS1* as well as the downregulated nodes *F2* in TCCSUPPi cells was carried out. Likewise, the upregulated node *RAC1* and downregulated node *TP53* obtained from EJ28Pi cells were subsequently investigated. The results showed that among the three hub genes (*RPL8, THBS1* and *F2*) obtained from TCCSUPPi cells, *RPL8* was significantly upregulated in bladder cancer cells as compared to normal bladder tissue (GSE3167; Figure 7 upper left; *p*= 6.36E-5) [24]. In a study undertaken by Kim, Kim [25], relative expression of *THBS1* and *F2* were slightly higher in bladder cancer than in the normal bladder tissue but the difference was not statistically significant (GSE13507; Figure 7 upper middle & upper right; *p*>0.05), suggesting that *RPL8, THBS1* and *F2* genes play a vital role in bladder cancer progression and metastasis. These genes may also be the reason why cancer cells are able to maintain normal growth and development even after NDV infection as seen in the case of the established persistent TCCSUPPi cells.

**Figure 7:**
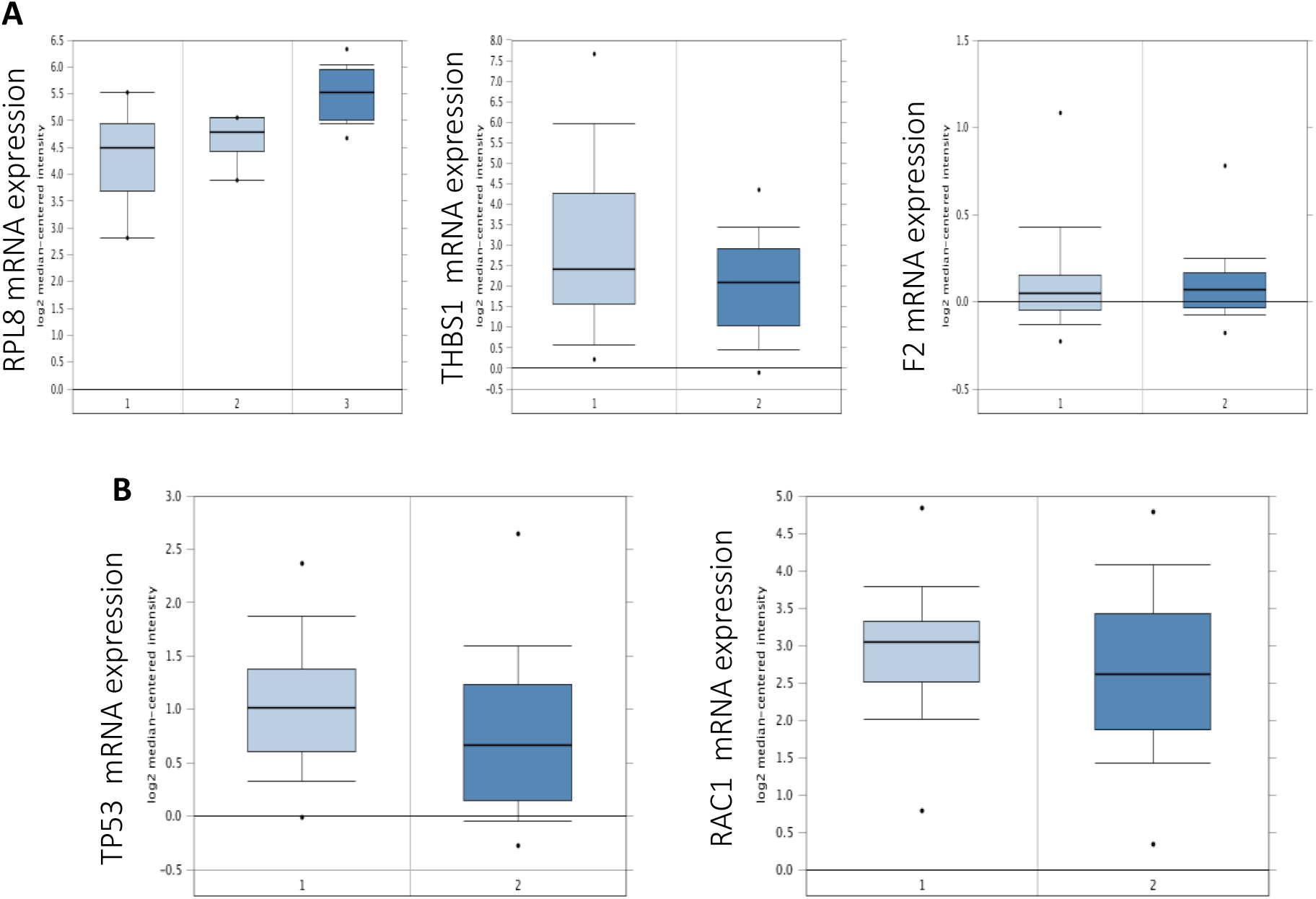
Expression of *RPL8, THBS1, F2, TP53,* and *RAC1* mRNA levels obtained from TCCSUPPi and EJ28Pi. A) Expression of *RPL8, THBS1* and *F2* mRNA levels obtained from TCCSUPPi in bladder cancer using Oncomine. B) Expression of *TP53* and *RAC1* mRNA levels obtained from EJ28Pi in bladder cancer using Oncomine. Left plot is normal bladder tissue while the right plot is bladder cancer tissue.

On the other hand, *TP53* and *RAC1* mRNA expression levels in bladder cancer is shown to be lower than in the normal bladder tissue (GSE13507; Figure 7 lower left & lower right; *p*>0.05). These data suggest that *TP53* and *RAC1* were downregulated in bladder cancer and may likely contribute to the development of NDV persistent infection in EJ28 cells noting that the *TP53* was found to be upregulated in the established persistent EJ28Pi cells.

## Discussion

Analysis of the RNA-Seq data revealed several essential pathways that were enriched by the DEGs identified to be involved in NDV persistent infection in bladder cancer cell lines. We then performed protein-protein interaction (PPI) analysis of the identified DEGs to unravel the connections between these pathways and the genes. It can be observed that there is functional connectivity in the biological processes. By modular analysis of the PPI network in persistent TCCSUPPi cells, *RPL8* node group was upregulated and implicated in the pathways of bladder cancer, malaria, mitophagy, p53 signaling, ECM-receptor interaction, TGF-beta signaling, phagosome, ribosome, focal adhesion and proteoglycans in cancer were clustered in the same module, suggesting that these genes co-function together to develop NDV persistent infection in TCCSUP bladder cancer cell line. *RPL8* is a protein coding gene being referred to as ribosomal protein L8 that is known to be associated with viral mRNA translation and rRNA processing pathways in both the nucleus and the cytosol[26]. In our study, *RPL8* gene was upregulated in the persistent TCCSUPPi cells. Dysregulation or misexpression of this gene family is associated with different types of cancer cells [27–30] and several other inherited genetic diseases[31]. This shows that *RPL8* is associated with viral infection and its role in establishing NDV persistent infection in TCCSUP bladder cancer cell line is obvious in this study.

On the other hand, the upregulated nodes found in persistent EJ28Pi cells includes *EP300, PTPN11,* and *RAC1* implicated in the pathways of renal cell carcinoma, viral carcinogenesis, proteoglycans in cancer, prostate cancer, insulin resistance, Ras signalling, circadian rhythm, neurotrophin signalling, cell cycle, aldosterone-regulated sodium reabsorption, FoxO signalling, microRNAs in cancer, Wnt signalling, influenza A, tight junction, viral myocarditis, Kaposi’s sarcoma-associated herpesvirus infection, PI3K-Akt signalling, epithelial cell signalling in helicobacter pylori infection, adipocytokine signalling, epstein-Barr virus infection, adherens junction, bacterial invasion of epithelial cells, cAMP signalling, HTLV-I infection, longevity regulating pathway, glucagon signalling pathway, leukocyte transendothelial migration, AMPK signalling pathway, pathways in cancer, natural killer cell mediated cytotoxicity and measles, were clustered in module 5, 6, and 7 (Figure 4.48), indicating that the genes in these cluster act together to facilitate the development of NDV persistent infection in EJ28 bladder cancer cell line. Our findings demonstrated that *EP300*, *PTPN11*, and *RAC1* genes were upregulated in NDV in persistent EJ28Pi cells. *EP300* gene is known to encode for adenovirus E1A-associated cellular p300 transcriptional co-activator protein that functions in regulations of transcription by remodeling chromatin[32–34]. It plays a vital role in cell proliferation, differentiation and epithelial cancer [35] as well as co-activate hypoxia-inducible factor 1 alpha, *HIF1A* to facilitate the expression of VEGF, a key player in inducing cellular hypoxia[36]. Recent study suggested that HAT domain mutation on *EP300* gene has a major impact on malignant progression and growth[37], suggesting that *EP300* gene may contribute to the survival and growth of EJ28 cells after being persistently infected with oncolytic NDV.

*PTPN11* also known as protein tyrosine phosphatase, none-receptor type 11 is an oncogene that has the potential to turn normal cells into cancerous cells when mutated. It encodes protein that is in the protein tyrosine phosphatase (PTP) family and provides key important functions in cell growth and differentiation. *PTPN11* gene is mutated in about 1.19% of all types of cancers including melanoma, carcinoma of the lung, malignant glioma and leukaemia[38]. This is consistent with our report, demonstrating that *PTPN11* gene is upregulated in NDV persistent EJ28Pi cells. In addition, *RAC1* RAS superfamily was also found to be upregulated in EJ28Pi cells. It functions by controlling cytoskeletal reorganization and cell growth. In line with our results, evidence has shown that the upregulation of *RAC1* is associated with lymphovascular invasion and lymph node metastasis of the urinary tract cancer[39]. Therefore, upregulation of *EP300, PTPN11*, and *RAC1* genes may play a role in the development of NDV persistent infection in EJ28 bladder cancer cell line via association with the above implicated pathways.

Protein-drug interaction network revealed drugs that may be used to disconnect the linkages between the major modules in the constructed PPI network for both the TCCSUPPi and EJ28Pi cells. We observed several drugs that are linked to the downregulated node *F2* in TCCSUPPi cells that includes lepirudin, bivalirudin, drotrecogin alfa, coagulation factor IX (recombinant), menadione, argatroban and proflavine etc. On the other hand, puromycin, anisomycin and alpha-hydroxyl-beta-phenyl-propionic acid were the drug targets for upregulated *RPL8* node group. One important drug among the connected drugs on *F2* node network is menadione, a biologically active vitamin K3 that promotes coagulation of blood and is reported to strongly inhibit cancerous growth especially the neoplastic cells through conversion to vitamin K2, and reduced mutagenicity in rapidly growing cells in a new born[40]. Other studies demonstrated synergistic effects of vitamin C and menadione (vitamin K3) alone with radiation therapy on killing of bladder cancer cells as well as other cancer types without adverse reactions for patients[41–43]. Anisomycin drug is an antibiotic that inhibits synthesis of protein via the activity of peptidyl transferase in ribosomes[44], it interacts with *RPL8* node in the TCCSUPPi PPI network. Studies have found that this drug kills a variety of cancer cells such as adenocarcinoma, colon and leukaemia through induction of apoptosis[45, 46]. Thus, anisomycin that interacts with *RPL8* as well as menadione and the remaining drugs that interacts with *F2* are drugs that can potentially reverse the mechanism employed by TCCSUP cells in acquiring NDV persistent infection thus may be used in combination with NDV to synergistically kill bladder cancer cells that initially resist NDV-mediated oncolysis.

The identified drugs that interact with upregulated rac family small GTPase 1 (*RAC1*) node in persistent EJ28Pi cells network include dextromethorphan and guanosine-5’-diphosphate. On the other hand, the drugs acetylsalicylic acid, AZD, and 1-(9-ethyl-9H-carbazol-3-yl)-N-methylmethanamine were found to interact with downregulated *TP53* node. Dextromethorphan belongs to a class of drugs known as morphinan that has sedative and stimulant properties, and is often used as cough suppressant. Report has it that dextromethorphan is capable of causing mild motor and cognitive impairment, paranoia and delusion[47]. It is unclear how dextromethorphan exerts its effects on bladder cancer cell lines. Thus, investigating the role of dextromethorphan on bladder cancer cell line is needed to understand its mechanism of action on cancer. We also found that guanosine-5’-diphosphate drug interacts with *RAC1* network. It has been reported that guanosine-5’-diphosphate in combination with other drug agents has significantly reduced the aggressive nature of breast cancers[48]. Another important drug identified to interact with *TP53* node in the network is AZD. This drug may potentially be used for bladder cancer treatment because AZD is already being used in clinical trials, testing against many different types of cancers including esophageal cancer [49], small cell lung cancer[50] and colorectal cancer[51]. Therefore, the drugs mentioned above may be used in combination with NDV to destroy the EJ28 bladder cancer cells and prevent them from further acquiring NDV persistent infection.

Our results further discovered that some upstream genes such as *RPL8* and *F2* were significantly upregulated in bladder cancer with the exception of *THBS1, TP53* and *RAC1* that were found to be less expressed according to the Oncomine-based expression analysis. Through PPI network analysis, we identified *RPL8* and *F2* as therapeutic drug targets which were obviously overexpressed in bladder cancer. The expression patterns indicate that *RPL8*, *F2, THBS1, TP53* and *RAC1* are differentially expressed in bladder cancer versus the normal bladder tissues and that the level of their expression varies depending on the nature of tissue.

## Conclusion

Our results for the first time analyzed and profiled the transcriptome of NDV persistent bladder cancer cell lines to construct protein-protein interaction (PPI) network. We observed that the upregulated *RPL8* group and downregulated *HSPA1A/HSPA4* group are functionally connected with one another based on the data from TCCSUPPi cells. Through *RPL8* - *HSPA1A/HSPA4* link, a considerable number of pathways were identified that includes bladder cancer, malaria, p53 signaling, ECM-receptor interaction and TGF-beta signaling were implicated which are indicative of the molecular pathways associated with NDV persistent infection in TCCSUP bladder cancer cell line. Similarly, we reported the functional connectivity between upregulated *EP300, IRS1*, *PTPN11*, and *RAC1* group and the downregulated *TP53*, *SP1*, *CCND1* and *XPO1* group based on the data from EJ28Pi cells. Through *IRS1*, *PTPN11*, and *RAC1* as well as *TP53*, *SP1*, *CCND1* and *XPO1* connections, several major pathways including renal cell carcinoma, viral carcinogenesis, cell cycle, FoxO signaling, pathway in cancer, NK cells mediated cytotoxicity, Ras signaling etc were implicated. This is an indication of the molecular pathways associated with NDV persistent infection in EJ28 bladder cancer cell line.

Furthermore, our data provide clues into identifying new drug targets that can synergistically act together with NDV to combat and destroy bladder cancer cells as well as reverse the ability to acquire NDV persistent infection. The results in this study therefore enhances our understanding of host response to NDV persistent infection by providing candidate genes and pathways that may be associated with persistent infection in bladder cancer cells. In addition, several proteins that are crucial to the development of persistent infection were identified through PPI network.

Finally, we recommend that the potential drugs to target the key nodes on the PPI network should be screened as this will help to identify new approaches for cancer treatment in combination with the oncolytic Newcastle disease virus. Overall, this work provides the molecular basis for further research to unravel the function of these novel pathways, candidate genes and specific proteins that seems to activate the phenotypes associated with NDV persistent infection in bladder cancer cells.

## Materials and Methods

### DEGs from persistently infected cells

The DEGs in this study were previously published in a preprint [13] and deposited in NCBI Gene Expression Omnibus (GEO) under accession number GSE140902 and released to the public (https://www.ncbi.nlm.nih.gov/geo/query/acc.cgi?acc=GSE140902). A total of 63 and 134 differentially expressed genes (DEGs) from persistent TCCSUPPi and EJ28Pi cells relative to their uninfected controls were obtained from the total RNA samples of these NDV persistently infected cells. Of 63 DEGs in TCCSUPPi cells, 25 genes were upregulated and 38 genes were downregulated. Whilst in EJ28Pi, 55 genes were upregulated and 79 genes were downregulated. These DEGs were enriched into pathways as reported in the paper and were used for the PPI network construct and analysis.

### PPI network analysis

DEGs obtained from the persistently infected cells were used to construct the PPI network using a multifunctional online software Network Analyst (https://www.networkanalyst.ca) [52, 53]. Path and module analyses as well as determination of the protein-drug interactions were carried using this software. PPI networks were generated for each of the persistently infected cell line by submitting the number of DEGs together with their Ensembl gene IDs and Log2 (fold change) expression to the NetworkAnalyst software. Search Tool for the Retrieval of Interacting Genes/Proteins (STRING) Interactome was used as the PPI database and the cut off confidence scores was set at 900 since the range of the scores are between 400 to 1000[22]. The seeds were mapped to the corresponding molecular interaction database and the subnetworks with more cluster of nodes were chosen and demonstrated as top nodes in the network.

### Nodal path exploration

Nodes that are functionally connected and linked in the generated PPI network was visualized using the path explorer function segment of the NetworkAnalyst software[52]. Paths of interest and the connections between nodes were selected and redesigned to enable further understanding of the cascades of event in the PPI network. The implicated nodes that are significantly enriched in the pathways associated with development of persistent infection were specifically marked in the connections.

### Exploring module in the PPI network

Using the same NetworkAnalyst software[52], a module explorer section of the application was used to identify the clustered of subnetworks that collectively function together in the PPI network through Walktrap Algorithm that uses walk based strategy to detect module. The degrees of internal (edges within a module) and external (edges connecting the nodes of a module with the rest of the graph) edges for determining the significance of the module was based on the Wilcoxon rank-sum and values of*p* <0.05 were considered significant. Then, the modules of interest were marked with different colours in the PPI network.

### Protein drug interaction

The upregulated and downregulated nodes of interest obtained from persistently infected TCCSUPPi (upregulated *RPL8* and downregulated *HSPA1A/HSPA4* groups) and EJ28Pi (upregulated *EP300, IRS1, PTPN11,* and *RAC1* and downregulated *TP53, SP1, CCND1* and *XPO1)* cells were further analyzed to identify their interactions with drugs. DrugBank database version 5.0 was used to match the these nodes against the panel of drug target to generate and collect the protein-drug interactions network information[54].

### Validation of hub genes

The candidate hub genes identified from hub module in the network were validated by an Oncomine database (http://www.oncomine.org/)[23]. Expression level of these hub genes in both the persistently infected cell lines were validated using Dyrskjøt, Kruhøffer [24] (GSE3167) and Kim, Kim [25] (GSE13507) gene expression data with defined thresholds values for fold change >2, gene rank top 10% and *p* <0.001. Data type was restricted to only mRNA and statistical significance was based on Student’s *t* test.

## Supporting information

Supplementary Tables_All

## Data availability

The authors declare that all the data in this manuscript are available. Raw and processed data were deposited in NCBI Gene Expression Omnibus (GEO) under accession number GSE140902 (https://www.ncbi.nlm.nih.gov/geo/query/acc.cgi?acc=GSE140902).

## Acknowledgement

This study was supported by the Ministry Energy, Science, Technology, Environment and Climate Change (MESTECC) Malaysia Flagship Fund, reference number: FP0514B0021- 2(DSTIN).

## Author Contributions

U.A, A.V., S.C.C., D.M.C., S.L.C., S.A., and Y.K., designed the study. U.A. performed the wet and dry lab works as well as analysed the data. U.A wrote the paper. All authors reviewed the manuscript.

