## Supplementary Tables_All for "Analysis of PPI networks of transcriptomic expression identifies hub genes associated with Newcastle disease virus persistent infection in bladder cancer"

**Supplementary information**

**Supplementary Table 1: List of significantly enriched pathways based on upregulated DEGs connecting the nodes in subnetwork 1 for TCCSUPPi.**

| S/No. | Pathway | Total | Hits | P.Value | FDR |
|---|---|---|---|---|---|
| 1 | Bladder cancer | 41 | 1 | 0.0158 | 1 |
| 2 | Malaria | 49 | 1 | 0.0189 | 1 |
| 3 | Mitophagy - animal | 65 | 1 | 0.025 | 1 |
| 4 | p53 signaling pathway | 72 | 1 | 0.0277 | 1 |
| 5 | ECM-receptor interaction | 82 | 1 | 0.0315 | 1 |
| 6 | TGF-beta signaling pathway | 92 | 1 | 0.0353 | 1 |
| 7 | Phagosome | 152 | 1 | 0.0578 | 1 |
| 8 | Ribosome | 153 | 1 | 0.0582 | 1 |
| 9 | Focal adhesion | 199 | 1 | 0.0752 | 1 |
| 10 | Proteoglycans in cancer | 201 | 1 | 0.0759 | 1 |

**Supplementary Table 2: List of significantly enriched pathways based on downregulated DEGs connecting the nodes in subnetwork 1 for TCCSUPPi.**

| S/N. | Pathway | Total | Hits | P.Value | FDR |
|---|---|---|---|---|---|
| 1 | Antigen processing and presentation | 77 | 3 | 3.76E-06 | 0.0012 |
| 2 | Protein processing in endoplasmic reticulum | 165 | 2 | 0.00264 | 0.419 |
| 3 | Prion diseases | 35 | 1 | 0.018 | 1 |
| 4 | Legionellosis | 55 | 1 | 0.0281 | 1 |
| 5 | Longevity regulating pathway - multiple species | 62 | 1 | 0.0317 | 1 |
| 6 | Complement and coagulation cascades | 79 | 1 | 0.0402 | 1 |
| 7 | Chagas disease (American trypanosomiasis) | 103 | 1 | 0.0522 | 1 |
| 8 | Toxoplasmosis | 113 | 1 | 0.0572 | 1 |
| 9 | Platelet activation | 124 | 1 | 0.0626 | 1 |
| 10 | Spliceosome | 134 | 1 | 0.0675 | 1 |

### Supplementary Table 3: List of drugs targets from TCCSUPPI protein-drugs

| S/N | ID | Label | Degree | Betweenness |
|---|---|---|---|---|
| 1 | P00734 | F2 | 103 | 5253 |
| 2 | DB00001 | Lepirudin | 1 | 0 |
| 3 | DB00006 | Bivalirudin | 1 | 0 |
| 4 | DB00055 | Drotrecogin alfa | 1 | 0 |
| 5 | DB00100 | Coagulation Factor IX (Recombinant) | 1 | 0 |
| 6 | DB00170 | Menadione | 1 | 0 |
| 7 | DB00278 | Argatroban | 1 | 0 |
| 8 | DB01123 | Proflavine | 1 | 0 |
| 9 | DB01766 | Beta-(2-Naphthyl)-Alanine | 1 | 0 |
| 10 | DB01767 | Hemi-Babim | 1 | 0 |
| 11 | DB02193 | 2-(2-Hydroxy-Phenyl)-1h-Benzoimidazole-5-Carboxamidine | 1 | 0 |
| 12 | DB02351 | Hirulog | 1 | 0 |
| 13 | DB02723 | 4-Oxo-2-Phenylmethanesulfonyl-Octahydro-Pyrrolo[1,2-a]Pyrazine-6-Carboxylic Acid [1-(N-Hydroxycarbamimidoyl)-Piperidin-4-Ylmethyl]-Amide | 1 | 0 |
| 14 | DB03136 | 4-Iodobenzo[B]Thiophene-2-Carboxamidine | 1 | 0 |
| 15 | DB03159 | CRA_8696 | 1 | 0 |
| 16 | DB03847 | Gamma-Carboxy-Glutamic Acid | 1 | 0 |
| 17 | DB03865 | 6-Chloro-2-(2-Hydroxy-Biphenyl-3-Yl)-1h-Indole-5-Carboxamidine | 1 | 0 |
| 18 | DB04136 | Lysophosphotidylserine | 1 | 0 |
| 19 | DB04591 | N-{2,2-DIFLUORO-2-[(2R)-PIPERIDIN-2-YL]ETHYL}-2-[2-(1H-1,2,4-TRIAZOL-1-YL)BENZYL][1,3]OXAZOLO[4,5-C]PYRIDIN-4-AMINE | 1 | 0 |
| 20 | DB04697 | TRANS-4-(GUANIDINOMETHYL)-CYCLOHEXANE-L-YL-D-3-CYCLOHEXYLALANYL-L-AZETIDINE-2-YL-D-TYROSINYL-L-HOMOARGININAMIDE | 1 | 0 |
| 21 | DB04722 | 2-(3-CHLORO-6-{[2,2-DIFLUORO-2-(1-OXIDOPYRIDIN-2-YL)ETHYL]AMINO}-1-OXIDOPYRIDIN-2-YL)-N-[1-(3-CHLOROPHENYL)ETHYL]ACETAMIDE | 1 | 0 |
| 22 | DB04771 | 1-GUANIDINO-4-(N-NITRO-BENZOYLAMINO-L-LEUCYL-L-PROLYLAMINO)BUTANE | 1 | 0 |
| 23 | DB04772 | 1-GUANIDINO-4-(N-PHENYLMETHANESULFONYL-L-LEUCYL-L-PROLYLAMINO)BUTANE | 1 | 0 |
| 24 | DB04786 | Suramin | 1 | 0 |
| 25 | DB04898 | Ximelagatran | 1 | 0 |
| 26 | DB05777 | ART-123 | 1 | 0 |
| 27 | DB06404 | C1 Esterase Inhibitor (Human) | 1 | 0 |
| 28 | DB06695 | Dabigatran etexilate | 1 | 0 |
| 29 | DB06838 | methyl L-phenylalaninate | 1 | 0 |
| 30 | DB06841 | D-phenylalanyl-N-[(1S)-4-{[amino(iminio)methyl]amino}-1-(chloroacetyl)butyl]-L-prolinamide | 1 | 0 |
| 31 | DB06845 | (S)-N-(4-carbamimidoylbenzyl)-1-(2-(cyclopentylamino)ethanoyl)pyrrolidine-2-carboxamide | 1 | 0 |
| 32 | DB06850 | (S)-N-(4-carbamimidoylbenzyl)-1-(2-(cyclohexylamino)ethanoyl)pyrrolidine-2-carboxamide | 1 | 0 |
| 33 | DB06853 | N-cycloheptylglycyl-N-(4-carbamimidoylbenzyl)-L-prolinamide | 1 | 0 |
| 34 | DB06854 | 2-(2-HYDROXY-BIPHENYL)-1H-BENZOIMIDAZOLE-5-CARBOXAMIDINE | 1 | 0 |
| 35 | DB06858 | N-cyclooctylglycyl-N-(4-carbamimidoylbenzyl)-L-prolinamide | 1 | 0 |
| 36 | DB06859 | N-ALLYL-5-AMIDINOAMINOOXY-PROPYLOXY-3-CHLORO-N-CYCLOPENTYLBENZAMIDE | 1 | 0 |
| **S/N** | **ID** | **Label** | **Degree** | **Betweenness** |

| S/N | ID | Label | Degree | Betweenness |
|---|---|---|---|---|
| 37 | DB06861 | 6-(2-HYDROXY-CYCLOPENTYL)-7-OXO-HEPTANAMIDINE | 1 | 0 |
| 38 | DB06865 | 6-CARBAMIMIDOYL-2-[2-HYDROXY-6-(4-HYDROXY-PHENYL)-INDAN-1-YL]-HEXANOIC ACID | 1 | 0 |
| 39 | DB06866 | 6-CARBAMIMIDOYL-2-[2-HYDROXY-5-(3-METHOXY-PHENYL)-INDAN-1-YL]-HEXANOIC ACID | 1 | 0 |
| 40 | DB06868 | N-(3-chlorobenzyl)-1-(4-methylpentanoyl)-L-prolinamide | 1 | 0 |
| 41 | DB06869 | 1-[2-AMINO-2-CYCLOHEXYL-ACETYL]-PYRROLIDINE-3-CARBOXYLIC ACID 5-CHLORO-2-(2-ETHYLCARBAMOYL-ETHOXY)-BENZYLAMIDE | 1 | 0 |
| 42 | DB06878 | 1-[(2R)-2-aminobutanoyl]-N-(3-chlorobenzyl)-L-prolinamide | 1 | 0 |
| 43 | DB06911 | D-leucyl-N-(3-chlorobenzyl)-L-prolinamide | 1 | 0 |
| 44 | DB06919 | D-phenylalanyl-N-(3-chlorobenzyl)-L-prolinamide | 1 | 0 |
| 45 | DB06929 | 1-butanoyl-N-(4-carbamimidoylbenzyl)-L-prolinamide | 1 | 0 |
| 46 | DB06936 | N-(4-carbamimidoylbenzyl)-1-(4-methylpentanoyl)-L-prolinamide | 1 | 0 |
| 47 | DB06942 | N-(4-carbamimidoylbenzyl)-1-(3-phenylpropanoyl)-L-prolinamide | 1 | 0 |
| 48 | DB06947 | 1-[(2R)-2-aminobutanoyl]-N-(4-carbamimidoylbenzyl)-L-prolinamide | 1 | 0 |
| 49 | DB06996 | D-leucyl-N-(4-carbamimidoylbenzyl)-L-prolinamide | 1 | 0 |
| 50 | DB07005 | D-phenylalanyl-N-{4-[amino(iminio)methyl]benzyl}-L-prolinamide | 1 | 0 |
| 51 | DB07016 | (3R)-8-(dioxidosulfanyl)-3-methyl-1,2,3,4-tetrahydroquinoline | 1 | 0 |
| 52 | DB07027 | D-phenylalanyl-N-(3-fluorobenzyl)-L-prolinamide | 1 | 0 |
| 53 | DB07083 | beta-phenyl-D-phenylalanyl-N-propyl-L-prolinamide | 1 | 0 |
| 54 | DB07088 | (S)-N-(4-carbamimidoylbenzyl)-1-(2-(cyclopentyloxy)ethanoyl)pyrrolidine-2-carboxamide | 1 | 0 |
| 55 | DB07091 | (S)-N-(4-carbamimidoylbenzyl)-1-(2-(cyclohexyloxy)ethanoyl)pyrrolidine-2-carboxamide | 1 | 0 |
| 56 | DB07095 | (S)-N-(4-carbamimidoylbenzyl)-1-(3-cyclopentylpropanoyl)pyrrolidine-2-carboxamide | 1 | 0 |
| 57 | DB07105 | 2-[2-(4-CHLORO-PHENYLSULFANYL)-ACETYLAMINO]-3-(4-GUANIDINO-PHENYL)-PROPIONAMIDE | 1 | 0 |
| 58 | DB07120 | N4-(N,N-DIPHENYLCARBAMOYL)-AMINOGUANIDINE | 1 | 0 |
| 59 | DB07128 | N7-BUTYL-N2-(5-CHLORO-2-METHYLPHENYL)-5-METHYL[1,2,4]TRIAZOLO[1,5-A]PYRIMIDINE-2,7-DIAMINE | 1 | 0 |
| 60 | DB07131 | (S)-N-(4-carbamimidoylbenzyl)-1-(3-cyclohexylpropanoyl)pyrrolidine-2-carboxamide | 1 | 0 |
| 61 | DB07133 | D-phenylalanyl-N-(3-methylbenzyl)-L-prolinamide | 1 | 0 |
| 62 | DB07143 | D-phenylalanyl-N-benzyl-L-prolinamide | 1 | 0 |
| 63 | DB07165 | N-(5-CHLORO-BENZO[B]THIOPHEN-3-YLMETHYL)-2-[6-CHLORO-OXO-3-(2-PYRIDIN-2-YL-ETHYLAMINO)-2H-PYRAZIN-1-YL]-ACETAMIDE | 1 | 0 |
| 64 | DB07190 | 3-cyclohexyl-D-alanyl-N-(3-chlorobenzyl)-L-prolinamide | 1 | 0 |
| 65 | DB07211 | (2R)-2-(5-CHLORO-2-THIENYL)-N-{(3S)-1-[(1S)-1-METHYL-2-MORPHOLIN-4-YL-2-OXOETHYL]-2-OXOPYRROLIDIN-3-YL}PROPENE-1-SULFONAMIDE | 1 | 0 |
| 66 | DB07277 | 2-(5-CHLORO-2-THIENYL)-N-{(3S)-1-[(1S)-1-METHYL-2-MORPHOLIN-4-YL-2-OXOETHYL]-2-OXOPYRROLIDIN-3-YL}ETHANESULFONAMIDE | 1 | 0 |
| 67 | DB07278 | 2-(5-CHLORO-2-THIENYL)-N-{(3S)-1-[(1S)-1-METHYL-2-MORPHOLIN-4-YL-2-OXOETHYL]-2-OXOPYRROLIDIN-3-YL}ETHENESULFONAMIDE | 1 | 0 |
| 68 | DB07279 | N-ETHYL-N-ISOPROPYL-3-METHYL-5-{[(2S)-2-(PYRIDIN-4-YLAMINO)PROPYL]OXY}BENZAMIDE | 1 | 0 |
| **S/N** | **ID** | **Label** | **Degree** | **Betweenness** |
| 69 | DB07353 | 4-(2,5-DIAMINO-5-HYDROXY-PENTYL)-PHENOL | 1 | 0 |

| | | | | |
|---|---|---|---|---|
| 70 | DB07366 | 2-[N'-(4-AMINO-BUTYL)-HYDRAZINOCARBONYL]-PYRROLIDINE-1-CARBOXYLIC ACID BENZYL ESTER | 1 | 0 |
| 71 | DB07376 | 5-(DIMETHYLAMINO)-1-NAPHTHALENESULFONIC ACID(DANSYL ACID) | 1 | 0 |
| 72 | DB07400 | 1-ETHOXYCARBONYL-D-PHE-PRO-2(4-AMINOBUTYL)HYDRAZINE | 1 | 0 |
| 73 | DB07440 | 4-TERT-BUTYLBENZENESULFONIC ACID | 1 | 0 |
| 74 | DB07461 | 3-AMINO-3-BENZYL-9-CARBOXAMIDE[4.3.0]BICYCLO-1,6-DIAZANONAN-2-ONE | 1 | 0 |
| 75 | DB07508 | 4-(5-BENZENESULFONYLAMINO-1-METHYL-1H-BENZOIMIDAZOL-2-YLMETHYL)-BENZAMIDINE | 1 | 0 |
| 76 | DB07515 | 1-(2-{[(6-AMINO-2-METHYLPYRIDIN-3-YL)METHYL]AMINO}ETHYL)-6-CHLORO-3-[(2,2-DIFLUORO-2-PYRIDIN-2-YLETHYL)AMINO]-1,4-DIHYDROPYRAZIN-2-OL | 1 | 0 |
| 77 | DB07521 | 6-CHLORO-1-(2-{[(5-CHLORO-1-BENZOTHIEN-3-YL)METHYL]AMINO}ETHYL)-3-[(2-PYRIDIN-2-YLETHYL)AMINO]-1,4-DIHYDROPYRAZIN-2-OL | 1 | 0 |
| 78 | DB07522 | N-[(2R,3S)-3-AMINO-2-HYDROXY-4-PHENYLBUTYL]NAPHTHALENE-2-SULFONAMIDE | 1 | 0 |
| 79 | DB07527 | N-[(2R,3S)-3-AMINO-2-HYDROXY-4-PHENYLBUTYL]-4-METHOXY-2,3,6-TRIMETHYLBENZENESULFONAMIDE | 1 | 0 |
| 80 | DB07548 | 2-(6-CHLORO-3-{[2,2-DIFLUORO-2-(2-PYRIDINYL)ETHYL]AMINO}-2-OXO-1(2H)-PYRAZINYL)-N-[(2-FLUORO-6-PYRIDINYL)METHYL]ACETAMIDE | 1 | 0 |
| 81 | DB07549 | 2-(6-CHLORO-3-{[2,2-DIFLUORO-2-(2-PYRIDINYL)ETHYL]AMINO}-2-OXO-1(2H)-PYRAZINYL)-N-[(2-FLUORO-3-METHYL-6-PYRIDINYL)METHYL]ACETAMIDE | 1 | 0 |
| 82 | DB07550 | 2-(6-CHLORO-3-{[2,2-DIFLUORO-2-(1-OXIDO-2-PYRIDINYL)ETHYL]AMINO}-2-OXO-1(2H)-PYRAZINYL)-N-[(2-FLUOROPHENYL)METHYL]ACETAMIDE | 1 | 0 |
| 83 | DB07639 | 3-(7-DIAMINOMETHYL-NAPHTHALEN-2-YL)-PROPIONIC ACID ETHYL ESTER | 1 | 0 |
| 84 | DB07658 | AC-(D)PHE-PRO-BOROLYS-OH | 1 | 0 |
| 85 | DB07659 | AC-(D)PHE-PRO-BOROHOMOLYS-OH | 1 | 0 |
| 86 | DB07660 | AC-(D)PHE-PRO-BOROHOMOORNITHINE-OH | 1 | 0 |
| 87 | DB07665 | N-[2-(carbamimidamidooxy)ethyl]-2-{6-cyano-3-[(2,2-difluoro-2-pyridin-2-ylethyl)amino]-2-fluorophenyl}acetamide | 1 | 0 |
| 88 | DB07718 | 3-(4-HYDROXY-PHENYL)PYRUVIC ACID | 1 | 0 |
| 89 | DB07741 | 4-(1R,3AS,4R,8AS,8BR)-[1-DIFLUOROMETHYL-2-(4-FLUOROBENZYL)-3-OXODECAHYDROPYRROLO[3,4-A]PYRROLIZIN-4-YL]BENZAMIDINE | 1 | 0 |
| 90 | DB07796 | (3ASR,4RS,8ASR,8BRS)-4-(2-(4-FLUOROBENZYL)-1,3-DIOXODEACAHYDROPYRROLO[3,4-A] PYRROLIZIN-4-YL)BENZAMIDINE | 1 | 0 |
| 91 | DB07809 | 4-({[4-(3-METHYLBENZOYL)PYRIDIN-2-YL]AMINO}METHYL)BENZENECARBOXIMIDAMIDE | 1 | 0 |
| 92 | DB07897 | 1-(HYDROXYMETHYLENEAMINO)-8-HYDROXY-OCTANE | 1 | 0 |
| 93 | DB07934 | [[CYCLOHEXANESULFONYL-GLYCYL]-3[PYRIDIN-4-YL-AMINOMETHYL]ALANYL]PIPERIDINE | 1 | 0 |
| 94 | DB07944 | N-{3-METHYL-5-[2-(PYRIDIN-4-YLAMINO)-ETHOXY]-PHENYL}-BENZENESULFONAMIDE | 1 | 0 |
| **S/N** | **ID** | **Label** | **Degree** | **Betweenness** |

| | | | | |
|---|---|---|---|---|
| 95 | DB07946 | N-[2-({[amino(imino)methyl]amino}oxy)ethyl]-2-{6-chloro-3-[(2,2-difluoro-2-phenylethyl)amino]-2-fluorophenyl}acetamide | 1 | 0 |
| 96 | DB08061 | 4-[3-(4-CHLOROPHENYL)-1H-PYRAZOL-5-YL]PIPERIDINE | 1 | 0 |
| 97 | DB08062 | 3-(4-CHLOROPHENYL)-5-(METHYLTHIO)-4H-1,2,4-TRIAZOLE | 1 | 0 |
| 98 | DB08152 | {(2S)-1-[N-(tert-butoxycarbonyl)glycyl]pyrrolidin-2-yl}methyl (3-chlorophenyl)acetate | 1 | 0 |
| 99 | DB08187 | METHYL-PHE-PRO-AMINO-CYCLOHEXYLGLYCINE | 1 | 0 |
| 100 | DB08254 | 2-NAPHTHALENESULFONIC ACID | 1 | 0 |
| 101 | DB08422 | [PHENYLALANINYL-PROLINYL]-[2-(PYRIDIN-4-YLAMINO)-ETHYL]-AMINE | 1 | 0 |
| 102 | DB08546 | 4-[(3AS,4R,7R,8AS,8BR)-2-(1,3-BENZODIOXOL-5-YLMETHYL)-7-HYDROXY-1,3-DIOXODECAHYDROPYRROLO[3,4-A]PYRROLIZIN-4-YL]BENZENECARBOXIMIDAMIDE | 1 | 0 |
| 103 | DB08624 | BENZOTHIAZOLE | 1 | 0 |

**Supplementary Table 4 : List of significantly enriched pathways based on upregulated DEGs connecting the nodes in subnetwork 1 for EJ28Pi.**

| S/No. | Pathway | Total | Hits | P.Value | FDR |
|---|---|---|---|---|---|
| 1 | Renal cell carcinoma | 69 | 4 | 2.52E-05 | 0.00802 |
| 2 | Viral carcinogenesis | 201 | 5 | 0.000127 | 0.0202 |
| 3 | Proteoglycans in cancer | 201 | 4 | 0.00154 | 0.127 |
| 4 | Prostate cancer | 97 | 3 | 0.00187 | 0.127 |
| 5 | Insulin resistance | 108 | 3 | 0.00254 | 0.127 |
| 6 | Ras signaling pathway | 232 | 4 | 0.00261 | 0.127 |
| 7 | Circadian rhythm | 31 | 2 | 0.00282 | 0.127 |
| 8 | Neurotrophin signaling pathway | 119 | 3 | 0.00334 | 0.127 |
| 9 | Cell cycle | 124 | 3 | 0.00375 | 0.127 |
| 10 | Aldosterone-regulated sodium reabsorption | 37 | 2 | 0.00401 | 0.127 |
| 11 | FoxO signaling pathway | 132 | 3 | 0.00447 | 0.129 |
| 12 | MicroRNAs in cancer | 299 | 4 | 0.00649 | 0.172 |
| 13 | Wnt signaling pathway | 158 | 3 | 0.00738 | 0.181 |
| 14 | Influenza A | 167 | 3 | 0.0086 | 0.191 |
| 15 | Tight junction | 170 | 3 | 0.00902 | 0.191 |
| 16 | Viral myocarditis | 59 | 2 | 0.00995 | 0.198 |
| 17 | Kaposi's sarcoma-associated herpesvirus infection | 186 | 3 | 0.0115 | 0.206 |
| 18 | PI3K-Akt signaling pathway | 354 | 4 | 0.0117 | 0.206 |
| 19 | Epithelial cell signaling in Helicobacter pylori infection | 68 | 2 | 0.0131 | 0.21 |
| 20 | Adipocytokine signaling pathway | 69 | 2 | 0.0134 | 0.21 |
| 21 | Epstein-Barr virus infection | 201 | 3 | 0.0142 | 0.21 |
| 22 | Adherens junction | 72 | 2 | 0.0146 | 0.21 |
| 23 | Bacterial invasion of epithelial cells | 74 | 2 | 0.0153 | 0.212 |
| 24 | cAMP signaling pathway | 212 | 3 | 0.0164 | 0.217 |
| 25 | HTLV-I infection | 219 | 3 | 0.0179 | 0.227 |
| 26 | Longevity regulating pathway | 89 | 2 | 0.0217 | 0.266 |
| 27 | Glucagon signaling pathway | 103 | 2 | 0.0285 | 0.336 |
| 28 | Leukocyte transendothelial migration | 112 | 2 | 0.0333 | 0.378 |
| 29 | AMPK signaling pathway | 120 | 2 | 0.0378 | 0.414 |
| 30 | Pathways in cancer | 530 | 4 | 0.0439 | 0.455 |
| 31 | Natural killer cell mediated cytotoxicity | 131 | 2 | 0.0443 | 0.455 |
| 32 | Measles | 138 | 2 | 0.0487 | 0.484 |

**Supplementary Table 5 : List of significantly enriched pathways based on downregulated DEGs connecting the nodes in subnetwork 1 for EJ28Pi.**

| S/No. | Pathway | Total | Hits | P.Value | FDR |
|---|---|---|---|---|---|
| 1 | Wnt signaling pathway | 158 | 5 | 6.63E-05 | 0.0211 |
| 2 | Proteoglycans in cancer | 201 | 5 | 0.000207 | 0.0327 |
| 3 | HTLV-I infection | 219 | 5 | 0.000308 | 0.0327 |
| 4 | Pathways in cancer | 530 | 7 | 0.000469 | 0.0373 |
| 5 | Breast cancer | 147 | 4 | 0.000701 | 0.0425 |
| 6 | Cellular senescence | 160 | 4 | 0.000963 | 0.0425 |
| 7 | Melanoma | 72 | 3 | 0.00105 | 0.0425 |
| 8 | Glioma | 75 | 3 | 0.00118 | 0.0425 |
| 9 | MAPK signaling pathway | 295 | 5 | 0.0012 | 0.0425 |
| 10 | Transcriptional misregulation in cancer | 186 | 4 | 0.00168 | 0.0536 |
| 11 | Focal adhesion | 199 | 4 | 0.00216 | 0.0623 |
| 12 | Prostate cancer | 97 | 3 | 0.00247 | 0.0623 |
| 13 | Endocrine resistance | 98 | 3 | 0.00255 | 0.0623 |
| 14 | Th17 cell differentiation | 107 | 3 | 0.00327 | 0.0743 |
| 15 | Thyroid hormone signaling pathway | 116 | 3 | 0.00411 | 0.0871 |
| 16 | Thyroid cancer | 37 | 2 | 0.00484 | 0.0962 |
| 17 | FoxO signaling pathway | 132 | 3 | 0.0059 | 0.105 |
| 18 | Bladder cancer | 41 | 2 | 0.00592 | 0.105 |
| 19 | Measles | 138 | 3 | 0.00667 | 0.112 |
| 20 | Oxytocin signaling pathway | 153 | 3 | 0.00886 | 0.127 |
| 21 | Hippo signaling pathway | 154 | 3 | 0.00902 | 0.127 |
| 22 | Amyotrophic lateral sclerosis (ALS) | 51 | 2 | 0.00905 | 0.127 |
| 23 | Hepatitis C | 155 | 3 | 0.00918 | 0.127 |
| 24 | Jak-STAT signaling pathway | 162 | 3 | 0.0104 | 0.134 |
| 25 | Hepatitis B | 163 | 3 | 0.0105 | 0.134 |
| 26 | Endometrial cancer | 58 | 2 | 0.0116 | 0.142 |
| 27 | Basal cell carcinoma | 63 | 2 | 0.0136 | 0.143 |
| 28 | Axon guidance | 181 | 3 | 0.014 | 0.143 |
| 29 | Mitophagy - animal | 65 | 2 | 0.0144 | 0.143 |
| 30 | Central carbon metabolism in cancer | 65 | 2 | 0.0144 | 0.143 |
| 31 | Inflammatory bowel disease (IBD) | 65 | 2 | 0.0144 | 0.143 |
| 32 | Non-small cell lung cancer | 66 | 2 | 0.0148 | 0.143 |
| 33 | Kaposi's sarcoma-associated herpesvirus infection | 186 | 3 | 0.015 | 0.143 |
| 34 | Long-term potentiation | 67 | 2 | 0.0153 | 0.143 |
| 35 | Amphetamine addiction | 68 | 2 | 0.0157 | 0.143 |
| 36 | PI3K-Akt signaling pathway | 354 | 4 | 0.0164 | 0.145 |
| 37 | p53 signaling pathway | 72 | 2 | 0.0175 | 0.15 |
| 38 | Platinum drug resistance | 73 | 2 | 0.018 | 0.15 |

| | | | | | |
|---|---|---|---|---|---|
| 39 | Pancreatic cancer | 75 | 2 | 0.0189 | 0.154 |
| 40 | Chronic myeloid leukemia | 76 | 2 | 0.0194 | 0.154 |
| 41 | Regulation of actin cytoskeleton | 214 | 3 | 0.0218 | 0.169 |
| 42 | Colorectal cancer | 86 | 2 | 0.0244 | 0.185 |
| 43 | Th1 and Th2 cell differentiation | 92 | 2 | 0.0277 | 0.204 |
| 44 | Small cell lung cancer | 93 | 2 | 0.0282 | 0.204 |
| 45 | Choline metabolism in cancer | 99 | 2 | 0.0317 | 0.224 |
| 46 | Neurotrophin signaling pathway | 119 | 2 | 0.0444 | 0.307 |
| 47 | Cell cycle | 124 | 2 | 0.0478 | 0.315 |
| 48 | Platelet activation | 124 | 2 | 0.0478 | 0.315 |
| 49 | Oocyte meiosis | 125 | 2 | 0.0485 | 0.315 |
